# Integrating deep learning with microfluidics for biophysical classification of sickle red blood cells

**DOI:** 10.1101/2020.07.01.181545

**Authors:** Niksa Praljak, Shamreen Iram, Utku Goreke, Gundeep Singh, Ailis Hill, Umut A. Gurkan, Michael Hinczewski

**Author notes:** These authors contributed equally to this work (Niksa Praljak, Shamreen Iram).

## Abstract

Sickle cell disease (SCD), a group of inherited blood disorders with significant morbidity and early mortality, affects a sizeable global demographic largely of African and Indian descent. It is manifested in a mutated form of hemoglobin that distorts the red blood cells into a characteristic sickle shape with altered biophysical properties. Sickle red blood cells (sRBCs) show heightened adhesive interactions with inflamed endothelium, triggering obstruction of blood vessels and painful vaso-occlusive crisis events. Numerous studies have reported microfluidic-assay-based disease monitoring tools which rely on quantifying adhesion characteristics of adhered sRBCs from high resolution channel images. The current workflow for analyzing images from these assays relies on manual cell counting and detailed morphological characterization by a specially trained worker, which is time and labor intensive. Moreover manual counts by different individuals are prone to artifacts due to user bias. We present here a standardized and reproducible image analysis workflow designed to tackle these issues, using a two part deep neural network architecture that works in tandem for automatic, fast and reliable segmentation and classification into subtypes of adhered cell images. Our training utilized an exhaustive data set of images generated by the SCD BioChip, a microfluidic assay which injects clinical whole blood samples into protein-functionalized microchannels, mimicking physiological conditions in the microvasculature. The automated image analysis performs robustly in comparison to human classification: accuracies were similar to or better than those of the trained personnel, while the overall analysis time was improved by two orders of magnitude.

## 1 Introduction

### 1.1 Background

Sickle cell disease (SCD) affects over 100,000 Americans and more than 4 million genetically predisposed individuals worldwide [1–4]. The affected demographic commonly draws on ancestral lineage from parts of Africa and India. The most common form of SCD is caused by a single mutation in the *β* globin gene, leading to the expression of an abnormal form of hemoglobin, HbS, in red blood cells (RBCs). Although SCD originates from a single deficit gene, there are many observed clinical sub-phenotypes associated with the disease. They are not mutually exclusive and some of the associated complications are seen to cluster together, suggesting independent genetic modifiers as their epidemiological underpinnings [1]. These sub-phenotypes are associated with different acute and/or chronic complications. Common acute complications include pain crises, acute chest syndrome, stroke and hepatic or splenic sequestration. More long term effects include chronic organ damage of the lungs, bones, heart, kidneys, brain, and reproductive organs [5]. The resultant heterogeneity among SCD patients belonging to different disease sub-phenotypes underlies the need for new methodologies to allow intensive patient specific evaluation and management in outpatient, inpatient and emergency department settings [6]. SCD also requires early diagnosis after birth and constant clinical monitoring through the life-span of the patient, the absence of which leaves them prone to reduced quality of life and premature mortality [7, 8].

The underlying biophysics of SCD hinges on associated complex dynamical phenomena playing out in the vascular flow environment. Mutated hemoglobin molecules expressed in affected sickle RBCs (sRBCs) have a tendency to polymerize in oxygen starved environments, forming long chains which distort the cell profile. The damaged cell membrane displays morphological sickling (distortion into a crescent shape) which dislocates the membrane molecules and leads to a stiffer membrane scaffolding. Consequently sRBCs are more adhesive and less deformable than healthy RBCs. This increased membrane rigidity, along with altered adhesion characteristics that heighten interactions with the endothelium and plasma, directly give rise to SCD’s key manifestation: recurring, painful vaso-occlusive crisis events triggered by sRBC aggregation and blood vessel clogging [4, 9, 10]. The problem thus lends itself very naturally towards exploration in a microfluidic or adhesion assay setup. An important line of investigation in such studies is the search for predictive indicators of disease severity in terms of biophysical rather than molecular markers [8, 11–13]. Microfluidic platforms used for evaluation of sRBC adhesion dynamics have the advantage of being able to directly use clinical whole blood taken from SCD patients [8, 9, 14–16]. This is a versatile laboratory setup that allows one to mimic the complex vascular environment, and realistically explore the multiple, interconnected factors at play. These devices are thus good candidate tools for batch quantitative analyses of the mechanisms occurring in micro-vasculature prior to and during crises, as well as for testing intervention mechanisms.

In this study, we focus on one particular microfluidic platform, the SCD Biochip [14, 17]—a customizable, in-vitro adhesion assay where the microchannels can be functionalized with various endothelial proteins, and integrated with a programmable syringe pump unit that can implement physiologically relevant flow conditions. The analysis of the data from clinical whole blood samples injected into the SCD Biochip and similar experimental approaches has been challenging, with a major bottleneck being manual counting and categorization of cells from complex phase contrast or bright field microscopic images. Manual quantification of these images is a rigorous, time consuming process and inherently reliant on skilled personnel. This makes it unsuitable for high throughput, operationally lightweight, easily replicable studies. For example, manual cell counting and classification into morphology based sub-groups using the SCD Biochip platform tends to take upwards of 3 hours per image for trained experts. The need for a reliable, fully automated image segmentation, classification, and analysis scheme is thus paramount.

Here we present a standardized and reproducible image analysis workflow that eliminates the need for user input and is capable of handling large amounts of data, by utilizing a fully automated, machine-learning-based framework that analyzes SCD BioChip assay images in a matter of minutes. Several earlier studies have explored deep learning approaches for automating SCD image analyses [18, 19]. These studies illustrated the power of deep learning to distinguish morphological details at different life stages of the cell. Here we attempt for the first time to use it in a context that closely mimics the micro-vasculature in vivo, with cells from whole blood adhering to endothelial proteins under flow conditions. Our goal is to design and implement a classification scheme that sorts the sRBCs on the basis of morphological differences that arise with progressive HbS polymerization and sickling—features which strongly correlate with changes to the sRBC’s bio-mechanical properties. Our processing pipeline has been set up to be of use as a high-throughput tool with detection, tracking, and counting capabilities that could be harnessed to assess visual bio-markers of disease severity. In the long term, this makes our workflow highly suitable for integration into comprehensive monitoring and diagnostic platforms designed for patient specific clinical interventions—a key component of emerging targeted and curative therapies.

### 1.2 Complexity of classification in whole blood imaging

While significant progress has been made in understanding SCD pathogenesis [4, 20], full characterization of the complex interplay of factors behind occlusion events, and designing appropriate advanced therapeutic interventions, remain significantly challenging. Part of the challenge lies in recreating the conditions of the complex vascular environment, which is the overall goal of the SCD BioChip around which our workflow is designed. Along with the ability to control aspects like applied shear forces and choice of channel proteins, the microfluidic BioChips work with clinical whole blood samples. This lets the experimental setup approximate in vivo conditions as closely as possible, at the cost of significantly increasing the complexity of the image processing problem. Here we describe the various categories of objects—of both cellular and extra-cellular origin—that show up in our channel images. The segmentation process must thus be able to not only identify the sRBCS, but also distinguish them from these other objects with a reliable degree of accuracy.

- **RBCs**: Healthy RBCs are easily identifiable from their circular shape with an apparent dimple arising from a top-down view of the bi-concave cell profile (Fig. 1A). Since the channels are functionalized with proteins showing preferential adherence for sRBCs, very few healthy RBCs show up in our images.
- **Adhered sRBCs**: SCD pathogenesis (progressive stages of HbS polymerization) causes diseased RBCs to undergo deformation of their cell profile, going from a round to a more elongated, spindle-like shape. Simultaneously, the bi-concavity starts distending outwards. Examples of such partially sickled cells are shown in Fig. 1B. Cells at a stage of advanced disease progression, accelerated in hypoxic environments, become highly needle-like in shape, and completely lose their concavity. Examples of such highly sickled cases are shown in Fig. 1C. These two categories of adhered sRBC also correlate with biomechanical characteristics of the cell membrane, and we will label them by their membrane deformability, as described in more detail in Section 1.3: *deformable* (partially sickled) and *non-deformable* (highly sickled) sRBCs.
- **White blood cells (WBCs)**: Laminin, our choice of functionalization protein for this study, has known sRBC binding capabilities, and shows little WBC adhesion. Thus our channel images exhibit WBCs with far less frequency relative to sRBCs. The WBCs can be identified from a regular, round shape and smooth appearance, with varying degrees of internal detail (Fig. 1D).
- **Non-functionally adhered objects:** The focal plane of the microscope objective in the experiments is set to the protein-functionalized bottom of the channel. Objects adhered to this surface are thus in focus. Due to the finite height of the channel, non-specifically adhered objects outside the focal plane—stuck to the PMMA coverslip on the channel (Fig. 1E, i-iii) or flowing by in motion (Fig. 1E, iv)—show up as out-of-focus objects. They exhibit characteristic diffraction rings or a blurred appearance.
- **Other unclassified objects:** Various categories of other objects can also appear in the images. Examples include platelet clusters (Fig. 1F, i), cellular debris from lysed cells (Fig. 1F, ii-iii), and dirt/dust (Fig. 1F, iv-v).

**Fig 1:**
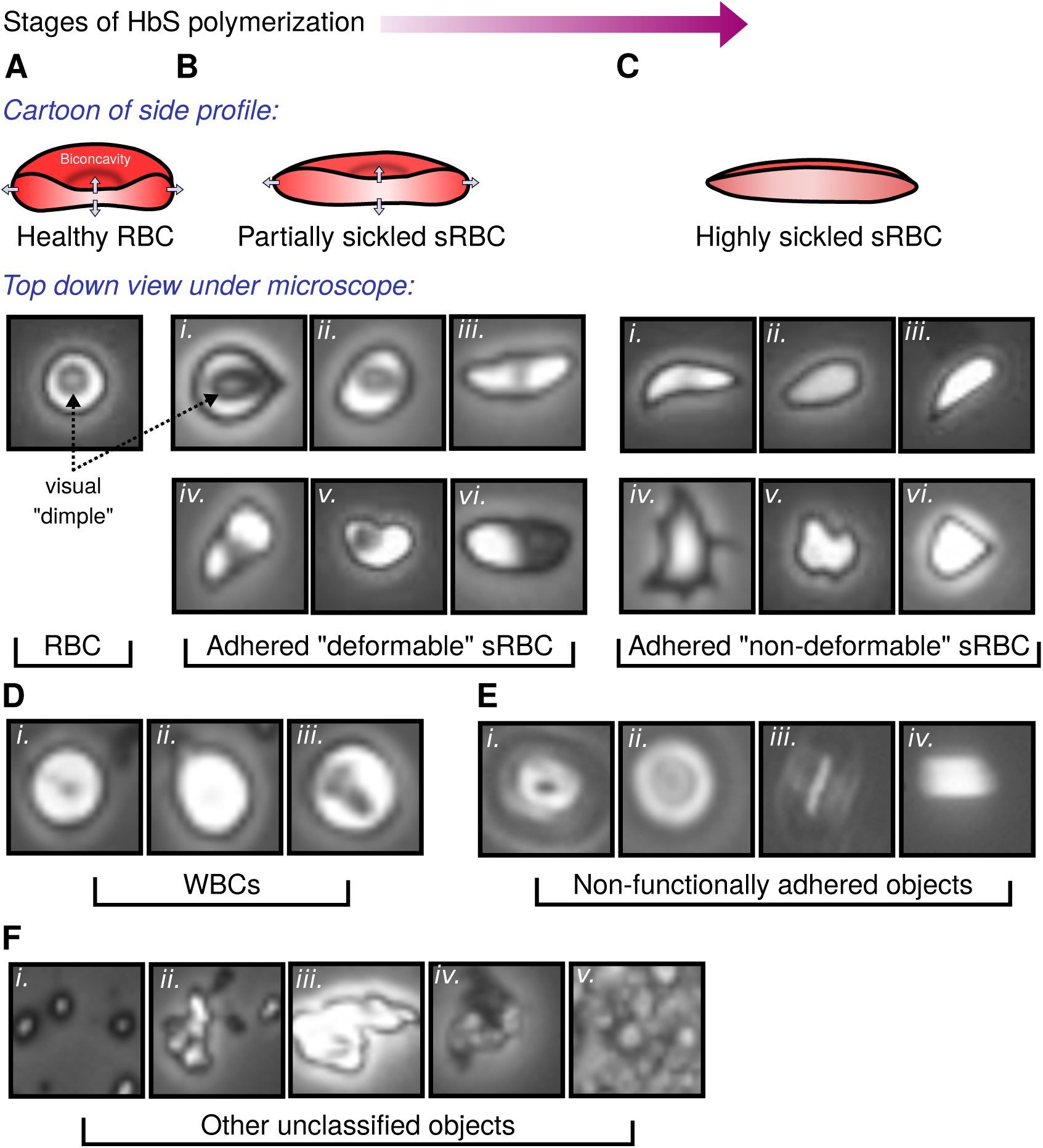
Object categories in our images: **(A-C)** The SCD pathogenetic pathway and changes undergone by the diseased RBC. **A:** A healthy RBC with biconcavity. The latter appears as a dimple viewed from the top. **B (i-iii):** Partially sickled sRBCs at increasing stages of sickling. The bi-concavity distends out to give a shallower dimple, and elongation in profile. This is the category we identify as *deformable* sRBC (see Section 1.3). **B (iv-vi):** Additional representative image variants of this category. **C (i-iii):** Highly sickled sRBCs. The dimple has completely disappeared and the shape is highly elongated. We classify these into our *non-deformable* category. **C (iv-vi):** More variants in the non-deformable category. Factors like local flow patterns, applied shear forces, and oxygen levels in the environment give rise to various shapes (teardrop, star-like, amorphous) for different sRBCs. **D:** White blood cells (WBCs). **E:** Non-functionally adhered objects. **F:** Other unclassified objects, like **(i)** platelet clusters, **(ii-iii)** lysed cells, **(iv-v)** dirt and dust. In our workflow types **D-F** are classified together in the *non-sRBC* category.

Along with these objects, the background itself can show considerable variation in luminosity and level of detail, depending on the sample and experimenter. A robust processing workflow should be able to deal with these challenges as well.

### 1.3 Establishing the biophysical basis for our classification problem

The observed range of heterogeneities in SCD clinical sub-phenotypes, stemming from the same monogenic underlying cause, remain ill understood [1, 21]. This lack of understanding sets the basis for the specific target problem that motivated our deep learning workflow. In a 2014 study from our group, Alapan *et al*. observed heterogeneity within sRBCs in terms of deformability and adhesion strength [9]—two biophysical characteristics that are key hallmarks of SCD pathogenesis. This observation revealed two new sub-classes with distinct morphologies, called deformable and non-deformable sRBCs (see Fig. 1B-C). The cells corresponding to the deformable class retain the RBC bi-concave detail, while the non-deformable sRBCs completely lose the bi-concave feature. This bi-concave feature for deformable sRBCs is visible to the naked eye in most ideal cases (see Fig. 1B i, ii, and v). However, based on the variety of configurations of the cell while adhered to the microchannel wall (see Fig. 1B iv and vi), in many cases detecting deformable sRBCs via human analysis can be complicated and inconsistent. These difficulties underline the importance of implementing deep learning models to quickly and consistently count and classify adhered cells.

As their name suggests, deformable sRBCs have relatively flexible shapes that can easily deform under a variety of physiologically relevant flow conditions. However, as RBCs fully progress through the sickling process, not only do the cells ultimately lose the concave dimple feature, but they also become stiffer. These so-called non-deformable sRBCs are readily distinguishable from deformable RBCs based on their sharper edges along with their missing dimples (see Fig. 1C). Below in Results Section 3.1 we demonstrate that these morphological differences correlate to significantly altered biomechanical properties in our experimental setup. Furthermore, in addition to their deformability characteristics, these two types of cells are also distinguishable in terms of their adhesion strength to endothelial and sub-endothelial proteins under fluid forces, making them potentially significant for understanding the biophysics of vaso-occlusive crises. In subsequent experiments that integrated a micro-gas exchanger with microfluidics, SCD heterogeneity is more dramatic under hypoxic conditions [22], a known precursor for the onset of these crises.

A wealth of information can be extracted by studying the morphological heterogeneity of sRBCs as a predictive indicator relevant to SCD pathogenesis and adhesion dynamics. Thus our automated deep learning workflow focuses on the above described SCD heterogeneity: counting sRBCs adhered to endothelial proteins in our microfluidic setup, and classifying these adhered cells into deformable and non-deformable types. Because the input consists of complex microscopy images of whole blood, the approach has to reliably disregard non-adhered sRBCs (see Fig. 1E) and other miscellaneous objects (see Fig. 1F).

## 2 Materials and methods

### 2.1 Details of experimental assay and image collection

RBC adhesion was measured using an in vitro microfluidic platform developed by our group, the SCD Biochip [14, 17]. The SCD Biochip is fabricated by lamination of a polymethylmethacrylate (PMMA) plate, custom laser-cut double-sided adhesive film which has a thickness of 50*µ*m (3M, Two Harbors, MN) and an UltraStick adhesion glass slide (VWR, Radnor, PA). 15 *µ*l of whole blood collected from patients diagnosed with SCD at University Hospitals, Cleveland, Ohio, was perfused into the microchannels functionalized with laminin (Sigma-Aldrich, St. Louis, MO). Laminin is a sub-endothelial protein with preferential adherence to sRBCs over healthy RBCs [23], allowing us to focus on sRBC characterization.

Images for the deep learning analysis were collected using the following protocol. Shear stress was kept at 0.1 Pa, mimicking the average physiological levels in post-capillary venules. After the non-adherent cells were removed by rinsing the microchannels, microfluidic images were taken by an Olympus IX83 inverted motorized microscope. Mosaic images were recorded and then stitched together by Olympus CellSense live-cell imaging and analysis software coupled with an QImaging ExiBlue Fluorescense Camera. An Olympus 10x/0.25 long working distance objective lens was used for imaging.

For the separate deformability analysis of Results Section 3.1, channel images were first captured under constant flow conditions of 10 *µ*L/min (which corresponds to a shear rate of about 100/s), and then subsequently captured again after the flow was turned off.

### 2.2 Overview of the image analysis workflow

We designed a bipartite network consisting of two individually trained neural networks that work in tandem to quantify our whole channel microfluidic image data. The workflow contains two phases of analysis that involve convolutional neural nets for cell segmentation/detection and classification of adhered sRBCs. We found this bipartite approach helpful in streamlining our workflow, trimming unnecessary operational bulk, and significantly improving robustness and performance metrics.

A schematic of the processing pipeline described here is shown in Fig. 2. Each phase of the pipeline has been built around a separate neural network. Since we are dealing with vast amounts of complex microscopy data that contains a plethora of cellular objects under fluid flow, we created Phase 1 to deal with object detection of adhered sRBCs exclusively. For Phase 2 (Fig. 2F) we focused on the biophysical classification of sRBCs into deformable and non-deformable types. After collecting microchannel images from the SCD BioChip, the workflow first implements Phase 1, which consists of a convolutional neural net with an architecture that downsamples and then upsamples the input image data into a segmentation mask (Fig. 2A-C). The downsampling portion of the network constructs and learns feature vectors as input data for the upsampling part of the neural network, allowing it to find segmentation masks for the original input images [24]. We also include crop and concatenations, motivated by the success of U-Net, a semantic segmentation network heavily used in the biomedical image segmentation community [25].

**Fig 2:**
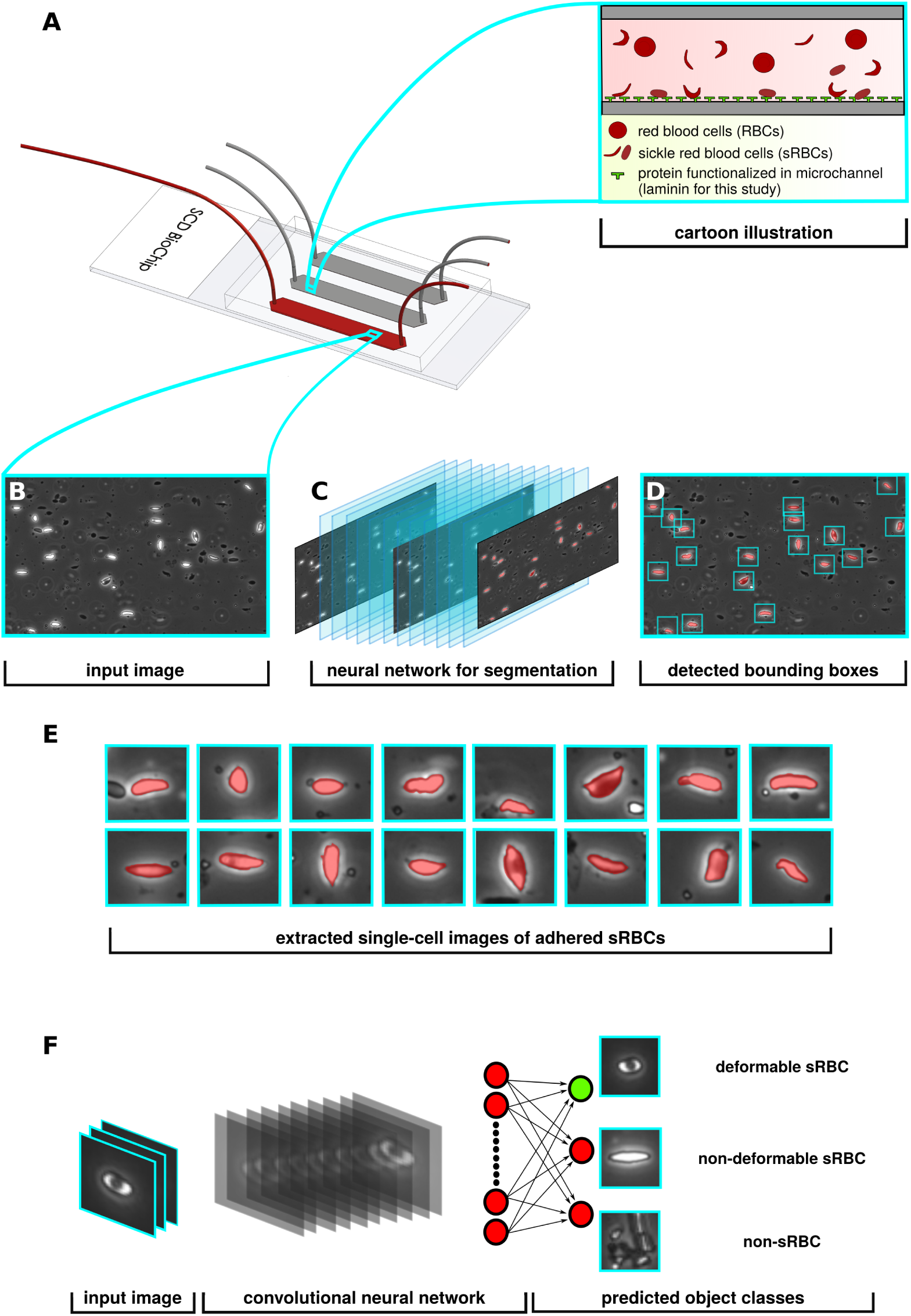
Overview of processing pipeline: **A:** SCD BioChip and cartoon illustration represents an in-vitro adhesion assay and adhesive dynamics of sRBCs within a mimicked microvasculature. **B:** Generated input image fed into the Phase I network. **C:** Phase I segmentation network predicts pixels belonging to adhered sRBCs, shaded red in the images. **D:** Drawing bounding boxes around segmented objects. **E:** Extracting adhered objects into individual images. **F:** The input layer of the Phase II classifier network receives an image from the Phase I detection network, then performs a series of convolutions and nonlinear activations to finally output class predictions.

Given an input image, the network learns to assign individual pixels to four categories (illustrated in Fig. 7A): background, deformable adhered sRBC, non-deformable adhered sRBC, and other. The other category largely involves detached or freely flowing cells (i.e. cells not attached to endothelial proteins along the channel wall, as seen in Fig. 1E), which are easily distinguishable from adhered cells. Training is done using a cross-entropy loss function penalizing wrong predictions of individual pixels.

**Fig 3:**
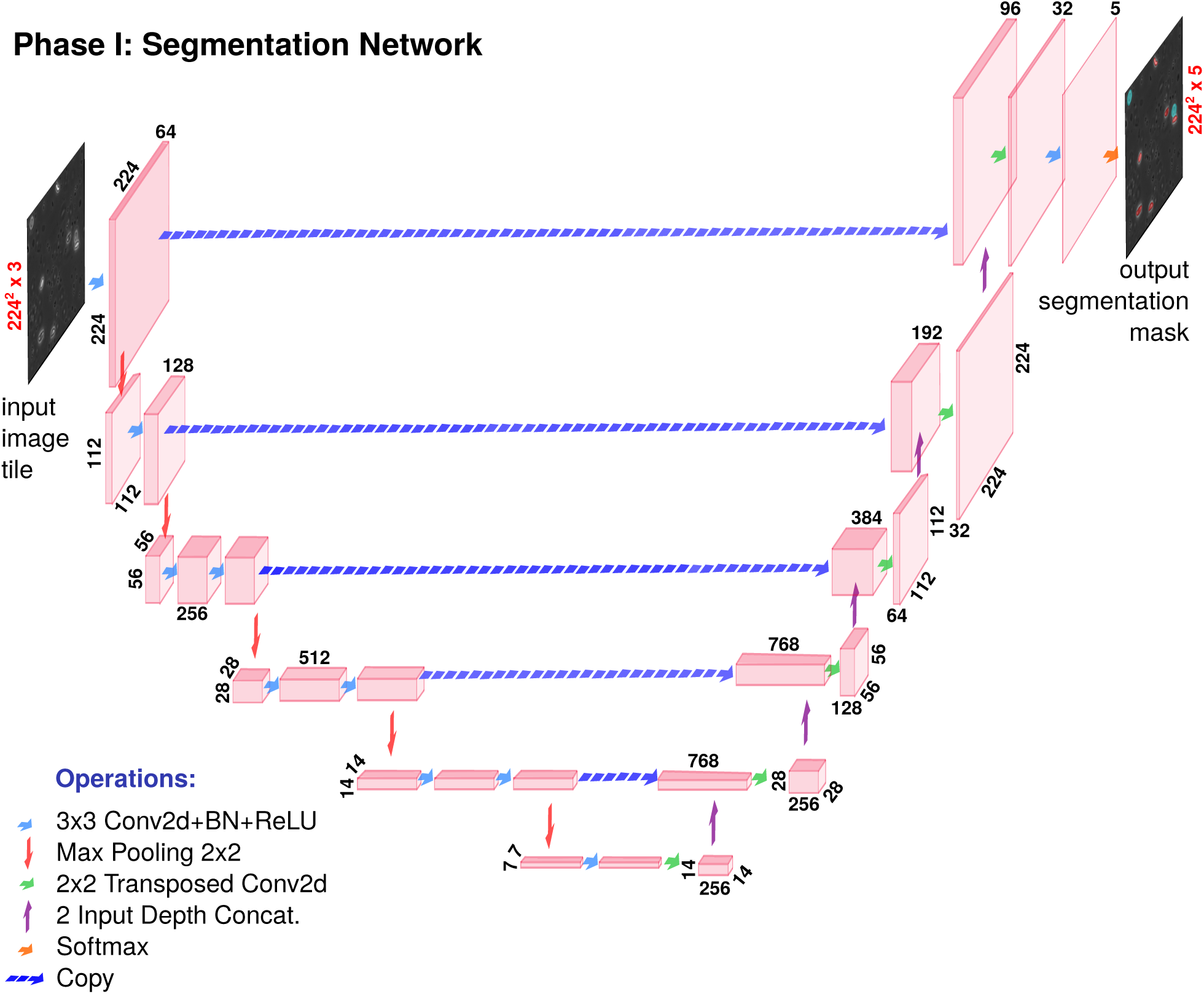
Phase I network: The architecture for our semantic segmentation convolutional neural network. The expanding and contracting arms making up the U shape are characteristic of U-Net-like networks. Dimension labels for each image volume are enumerated every time there is a change during a convolutional operation.

**Fig 4:**
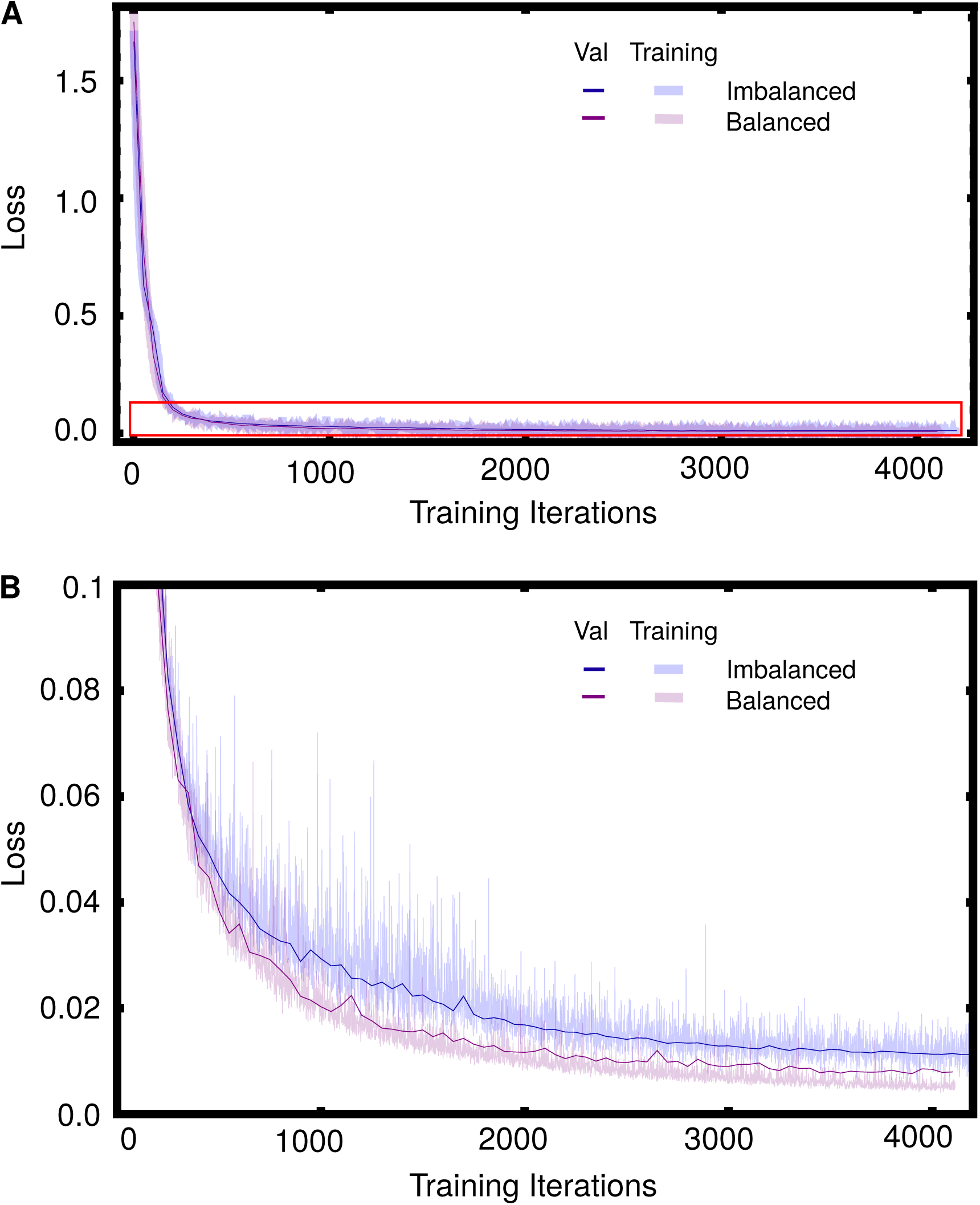
Phase I training and validation history: **(A)** Loss vs. training iterations for a network with no class balance (blue), and one with pre-training on a class-balanced data set (purple). Training and validation loss curves are shown in light-thick and dark-thin curves respectively. Training loss was evaluated at every iteration while validation loss was computed every 50 iterations. **(B)** shows a close up view of the loss ≤ 0.1 region of **(A)** (boxed in red).

**Fig 5:**
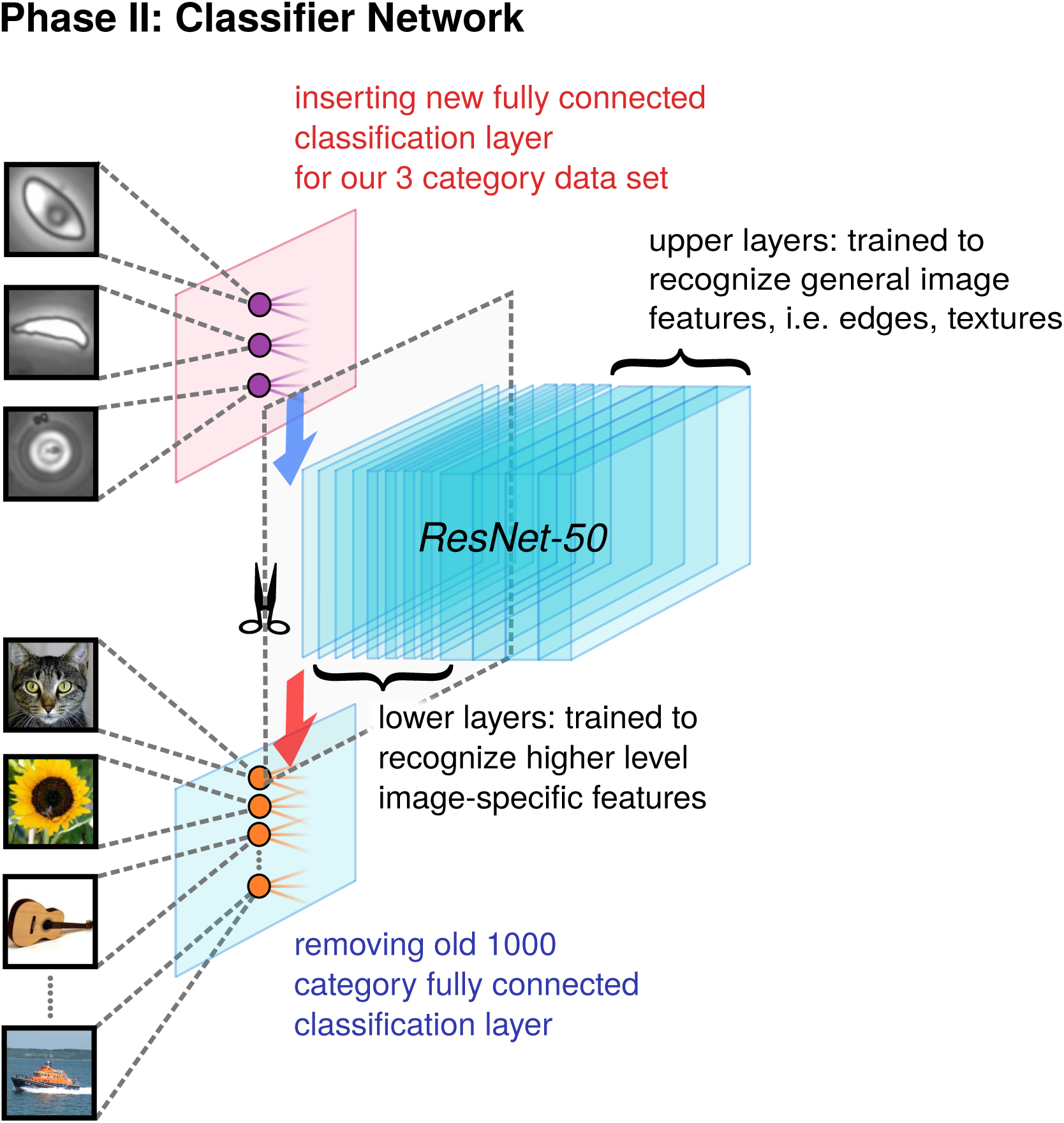
Phase II network: A schematic of the transfer learning workflow used to train our classifier network. We employ the ResNet-50 architecture and start with weights pre-trained on the 1000 category reduced ImageNet ILSRVC database. The final fully connected learnable classification layer is swapped out for a 3 class classification layer suited to our problem.

**Fig 6:**
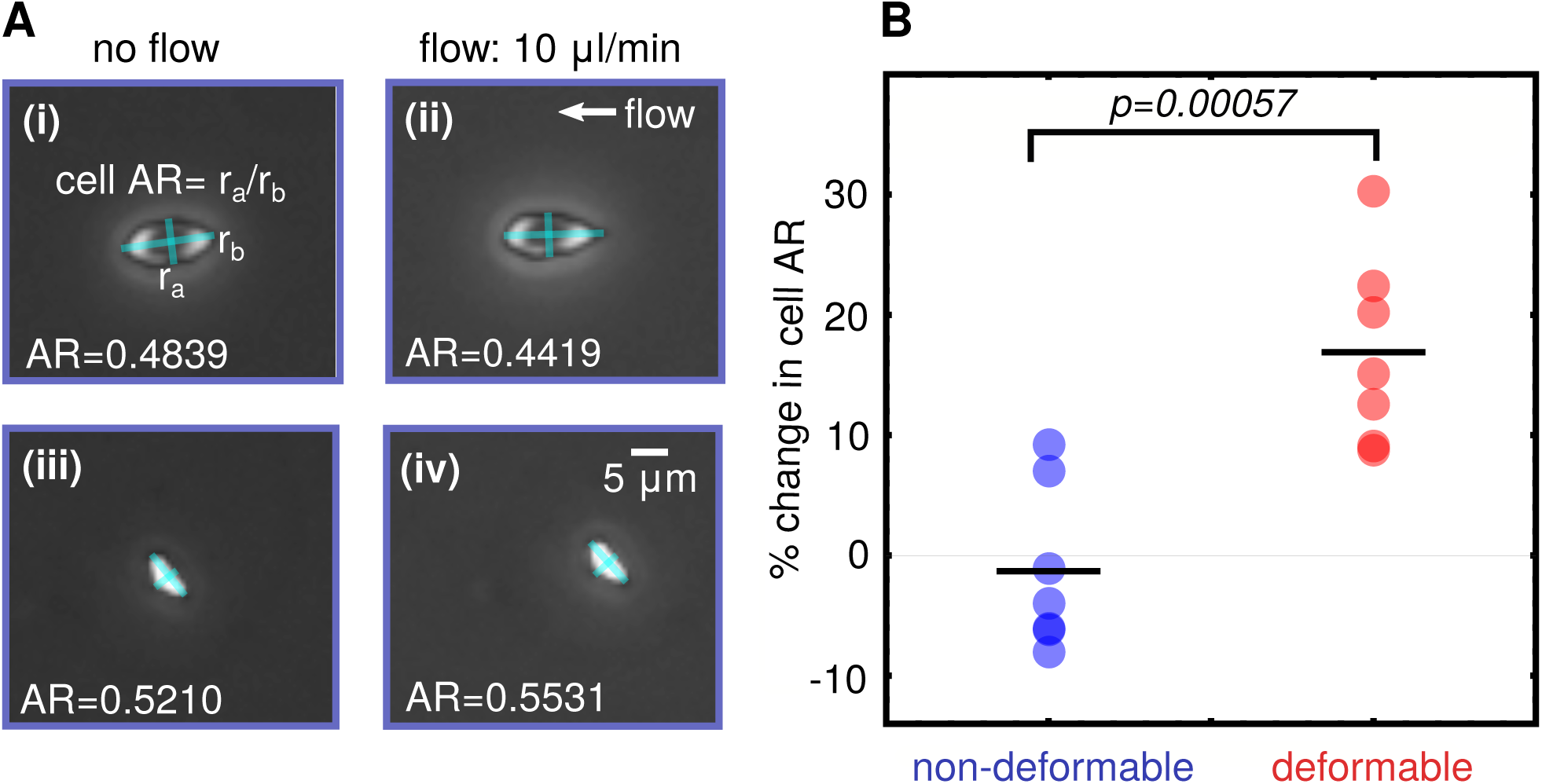
Cell deformability analysis: **A:** Schematic for estimation of change in cell aspect ratio (AR) between flow and no flow conditions. **(i-ii)** show a deformable type cell, and **(iii-iv)** a nondeformable. **B:** Mapping deformability to morphology: Cells visually identified as the deformable morphological subtype show significantly higher percentage change in cell AR between flow and no flow conditions compared to the non-deformable subtype.

**Fig 7:**
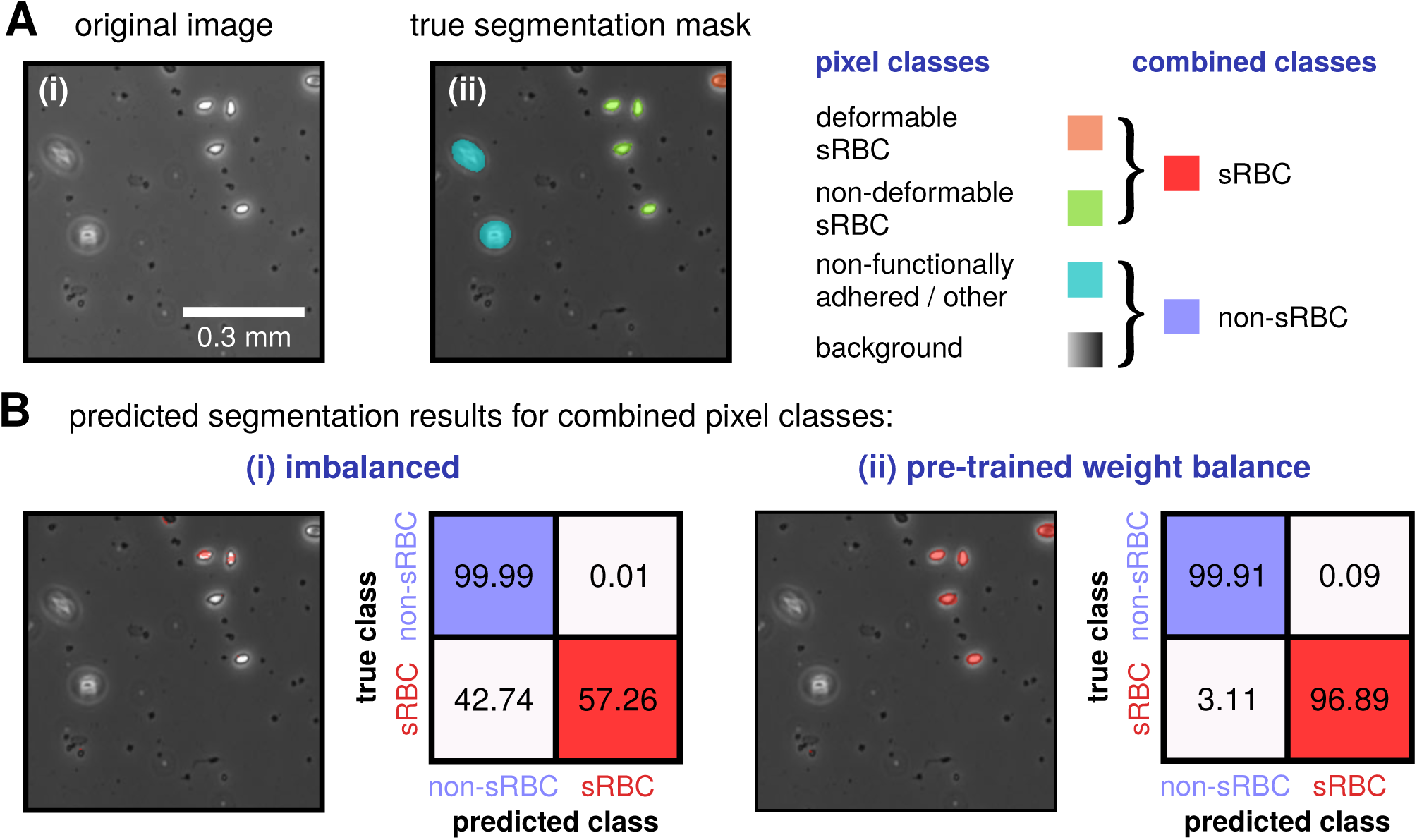
**(A)** Example of an input image tile for the Phase I network, along with the manually labeled segmentation mask assigning each pixel in the image to the classes listed on the right. **(B)** Given the predominance of background pixels in the image tiles, we use a pre-trained class balancing method for our Phase I network training. The results of this approach (panel ii) are compared against the case without any balancing (panel i). We show the segmentation mask produced by each network for the sample tile from **A**, with pixels predicted to be in the combined class sRBC colored red, while those predicted to be non-sRBC left uncolored. Comparison to the true segmentation mask shows that the pre-trained network does a much better job of distinguishing sRBC from non-sRBC pixels. This is quantified in the two confusion matrices from each network’s test run, expressed in terms of percentages. The matrix for the balanced network exhibits large diagonal elements (indicating class-wise prediction accuracy), and negligible off-diagonals (indicating low mis-classification).

The four-class pixel labeling scheme we have used for our Phase I segmentation network is a first step toward capturing the complexity of features of in our images. Ultimately however we are interested in classifying entire objects rather than individual pixels. Because the network might assign pixels from different classes to the same cell, there can be ambiguity in making definitive object classifications based on pixel labels (i.e. how to classify a cell with a comparable number of pixels assigned to the deformable and non-deformable sRBC classes). Thus further refinement is necessary, motivating the introduction of our Phase II network. To understand this issue in more detail, let us define two broader pixel label categories: *sRBC*, which is the combination of the two adhered sRBC pixel classes, and *non-sRBC*, which consists of the background / other classes. In Fig. 2D-E the pixels predicted to be sRBC by the network are shaded red, while the non-sRBC pixels are uncolored. As detailed in Section 2.3, with the appropriate training approach (in our case class weight balancing using transfer learning), the Phase I network does an excellent job distinguishing between these two broader classes. It is not as accurate in making the finer distinction between deformable and non-deformable adhered sRBCs. However we can at least rely on the Phase I network to accurately identify clusters of pixels as adhered sRBCs. The algorithm then computes 32 × 32 pixel bounding boxes around such clusters, each box centered around the cluster centroid (Fig. 2D). The size of the box takes into account the average size of the sRBCs in our channel images at 10x magnification, so that one box typically contains an entire cell. These boxes then form a set images (Fig. 2E) that are the input for our Phase II network.

In Phase II (Section 2.4), the images are run through a convolutional neural net for biophysical classification. The neural network performs a series of convolutions and filters, which ultimately classifies the image as a deformable sRBC, non-deformable sRBC or non-sRBC (Fig. 2F). If the Phase I analysis was entirely error-free, there would of course be no input images in Phase II corresponding to non-sRBC objects. But we include this category to filter out the rare mistakes made by the Phase I analysis, further enhancing the accuracy of the results at the completion of Phase II. Our dataset for Phase II consisted of a library of 6,863 manually classified cell images. Since this is a modestly-sized dataset for training an image classification network, we decided to implement transfer learning, an approach that can enhance training in cases with limited data [26]. We found that the deep residual network called ResNet-50 [27], pretrained on ImageNet [28], worked well in learning morphological features for our biophysical classification task. We also conducted a *k*-fold cross-validation protocol to estimate the accuracy of our machine learning model on test data. The details of the architectural design, data set pre-processing and preparation, progress checkpoints, and evaluation metrics for each network phase are presented in the next two sections.

### 2.3 Phase I: Detecting adhered sRBCs

This section is ordered as follows. First, we present the details of our neural network for semantic segmentation with an architecture inspired by SegNet [24] and U-Net [25]. Then we demonstrate an issue with imbalanced pixel classes during training and how we overcame this issue using a transfer learning approach. This approach utilizes weights transferred from a network trained on a more balanced data set [29]. The latter contains image instances of individual cells to jump-start the learning process. Finally, we illustrate overall performance of the network to detect adhered sRBC cells by presenting multiple relevant evaluation metrics.

#### Preprocessing of microchannel images and preparation of the data set

Before we implement the neural network for segmentation and detection, we record mosaic images of a single whole channel and stitch each image together, leading to a larger image with pixel dimensions 15,000 × 5,250. We then split the raw whole channel image into 1,000 equally-sized tiles by dividing the rectangular image with 50 vertical and 20 horizontal partitions, leading to tiles with pixel dimensions 300 × 262.

For our optimal architecture, the network has an input layer with size 224 × 224 × 3, with the first two dimensions representing height and width in pixels, and the last dimension representing three channels. Though our tile images were all grayscale, their format varied depending on the experimental procedure for recording the photo, with some having three channels and some just one channel. In the latter case we copy the first channel and then concatenate the copied channel two more times, creating images with three-channel depth. We then resize the width and height of the tile from 300 × 262 × 3 to 224 × 224 × 3 with a bicubic interpolation function, to match the input specfications of the network, and apply zero-centered normalization.

Since we are using supervised learning, we require that the data set be manually labeled beforehand. This was accomplished using the Image Labeler app in Matlab R2019a. As described above, each pixel is assigned to one of four labels: background, deformable sRBC, non-deformable sRBC, and other. Examples of an original tile and its labeled counterpart are shown in Fig. 7(i)-(ii).

#### Phase I network architecture

Our Phase I network implements five convolutional blocks that contain filters, nonlinear activation functions (ReLU), batch normalization, down (max), and up (transpose) sampling layers. Altogether, these layers sum to a total of 61 individual layers that take images as input and classify individual pixels into the four label categories to generate a segmentation mask. The full details of the network architecture are shown in Fig. 3.

#### Overcoming imbalanced classes by transfer learning

A common challenge in training semantic segmentation models is class imbalanced data [30]. A class imbalance occurs when the frequency of occurrence of one or more classes characterizing the data set differs significantly in representation, usually by several orders of magnitude, from instances of the other classes. This problem results in poor network performance in labeling the minority classes, a significant challenge for biomedical image segmentation in which frequently the minority class is the one under focus. A typical example is in pathologies such as inflammatory tissues or cancer lesions, where the aberrant tissue patch or lesion is much smaller in size compared to the whole image. This issue leads to reduced capacity for learning features that correlate to the lesions. For our microchannel images, the background far outstrips the adhered sRBCs in representation, heavily skewing the data set. In the absence of balancing, since our cross-entropy loss sums over all the pixels in an image, we find the network significantly misclassifies adhered sRBC pixels, in some cases completely ignoring them. Since our interest lies in accurately identifying the adhered cells, it is imperative to address this imbalance and improve accuracy for these minority classes.

We explored and tested a transfer learning-oriented method to overcome class imbalances within our pixel-labeled data. In place of a standard training procedure starting with starting weights drawn from a normal distribution, we transferred weights from a pre-trained network. The pre-training involves a more inherently class balanced data set: 2,295 manually extracted images of deformable and non-deformable adhered sRBCs, as well as non-sRBC objects like out-of-focus cells. Unlike the set of 1,000 tiles described earlier, which typically contain multiple cells per tile, in this case we use single cell images, with bounding boxes of 32 × 32. Because our Phase I network architecture has a fixed input layer size, we then resize these images to 224 × 224 with bicubic interpolation. The details of the two Phase I data sets (single-cell images and larger tiles) are summarized in Table 1. The single cell images used in this pre-training had a much lower fraction of background pixels relative to the tiles. Thus by pre-training the network on these images, the hope is that this biases the network to classify non-background pixels more accurately during the subsequent training on the tiles.

**Table 1:** Details of data sets used both network phases. At each stage 75% of the data is used for training the network, 10% is used for validation, and 15% for testing.

| Phase | Dataset |
| --- | --- |
| I | Initial: 2,295 pixel-labeled single-cell images (each $32 \times 32$ pixels)<br>Final: 1,000 pixel-labeled tiles (each $224 \times 224$ pixels) |
| II | 6,863 single-cell images (each $32 \times 32$ pixels) representing:<br>3,362 deformable sRBC, 1,449 non-deformable sRBC, 2,052 non-sRBC |

In both the pre-training stage and the training on tiles, 75% of the respective data sets are used to train the network, with the remainder reserved for validation (10%) and testing (15%). For the tile stage, to prevent overfitting we inspected the training and validation loss progress and manually aborted the training process after the network reached 21 epochs, approximately 25 mins (see Fig. 4). Section 2.3 lists hardware implementation details.

### 2.4 Phase II: Classification into morphological subtypes

#### Network structure and data set preparation

The structure of our Phase II cell classifier network was adapted from ResNet-50, the very deep residual neural network [27]. Residual neural networks implement skip connections in the hopes of avoiding vanishing gradients. Our implementation of ResNet-50 is pre-trained on the reduced ImageNet ILSVRC database, consisting of over 1 million training images and belonging to 1000 different classes [28].

As mentioned earlier, the input images for Phase II are 32 × 32 pixel images corresponding to single cells. Ideally these are all adhered sRBCs, but there is a tiny subset of non-sRBC objects, a source of error that the Phase II network is designed to mitigate. The details of constructing these single-cell images are as follows. Starting with the four-class segmentation mask generated at the end of Phase I, we binarize the pixels in these images according to our two combined pixel classes (Fig. 7B) by assigning 1 to sRBC pixels and 0 to non-sRBC pixels. We delete any small objects that form connected clusters of 1 pixels where the cluster size is smaller than 60. This threshold allows us to remove debris from the images, while being small enough relative to the range of sRBC cell sizes to preserve clusters that are actually sRBCs. We compute the centroids of the remaining clusters, ideally corresponding to sRBC cells, and extract 32 × 32 pixel bounding boxes centered at each cluster centroid (see Fig. 2 D-E). Before we input these extracted cell images into the Phase II neural network for biophysical classification, we resize the image from 32 × 32 × 3 to the corresponding ResNet50 input layer size of 224 × 224 × 3, and apply zero-centered normalization.

The training set for our supervised learning in Phase II consists of 6,863 single-cell images in three object categories: deformable sRBC (3,362 images), non-deformable sRBC (1,449 images), and non-sRBC (2,052 images). Examples of these images are shown Fig. 9A. In terms of curating our data set, we initially started with a batch of individual objects that were manually extracted from a large set of channel images displaying different luminescence and granularity features that covered the broad spectrum of sample and experimental variance (see Fig. 1A). However, after we completed our Phase I network, we expanded the data set to include the single-cell images generated by Phase I, though we manually verified the labels to correct any errors. Our data set also covers different physiological conditions like normoxia and hypoxia, which allows the resulting image processing pipeline to handle data from a wide range of SCD assays.

**Fig 8:**
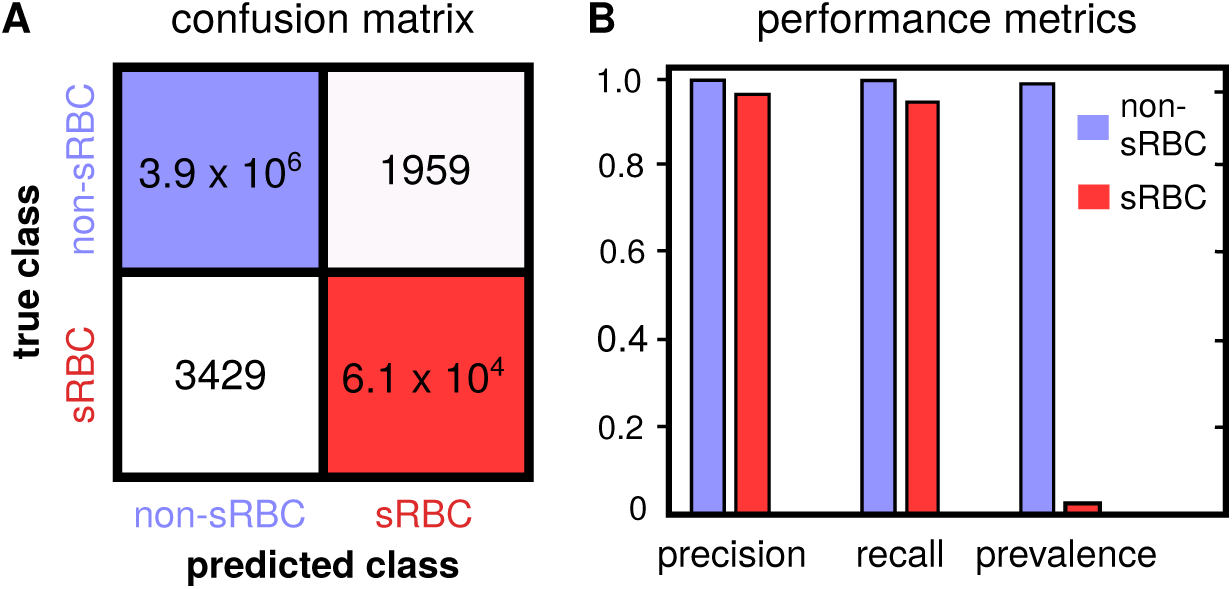
Phase I network performance metrics: **(A)** The customary confusion matrix in terms of pixel label counts rather than percentages of the predictions on the test data. **(B)** The precision, recall, and prevalence metrics, representing the overall performance of the network. The high values of precision and recall for the non-sRBC and sRBC categories testify to the ability of the network to correctly detect pixels belonging to sRBCs within large microchannel images. Prevalence illustrates the imbalance of non-sRBC vs. sRBC pixels in our data set.

**Fig 9:**
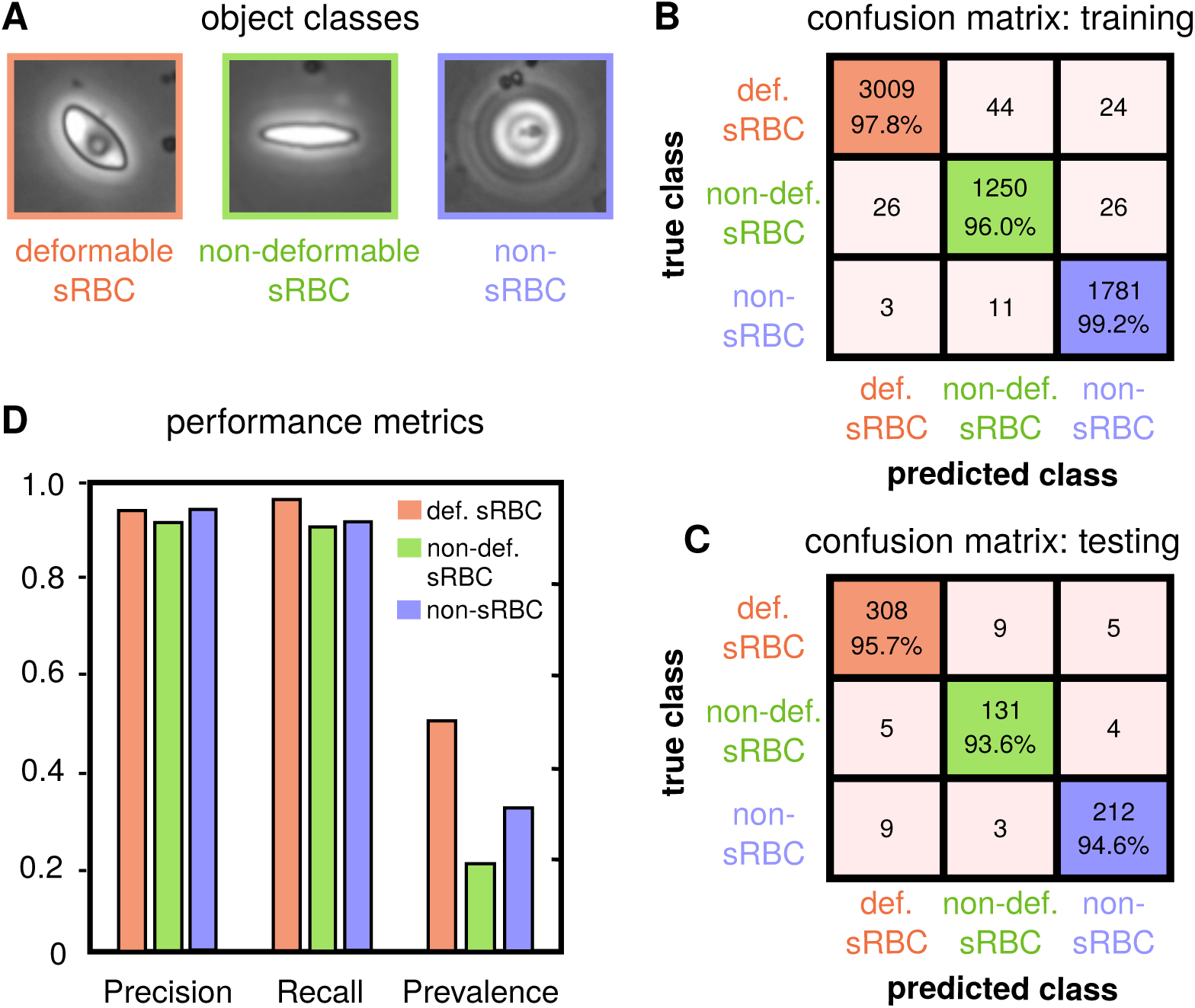
Phase II network performance metrics: **(A)** Representative examples of single-cell images for each classifier category, the input for Phase II. **(B-C)** Confusion matrices giving classifier performance accuracy on the training and testing sets (results shown for fold 1). Matrix elements are the numbers of each case in the data set, with percentages shown additionally along the diagonals. **(D)** Precision, recall and prevalence metrics for the network.

The overall data set size is still relatively small for our complex classification task, which requires learning subtle morphological features in cells of various sizes, shapes, and orientations. We thus choose to utilize a transfer learning framework [26]: rather than initializing the network with randomly chosen weights, we start with weights pre-trained on ImageNet, allowing us to achieve a higher starting accuracy on our own data set, faster convergence, and better asymptotic accuracy. To tailor the network for our purposes, we swapped out the final fully-connected layer of ResNet-50, which has output size 1000, to a layer with three output neurons corresponding to the three object classes (Fig. 5).

#### Phase II training details

The data set was split randomly into 80% training and 20% testing subsets, and the network was trained with maximum epoch number 10 and minibatch size 64. Each training session had 850 iterations, and thus 85 iterations per epoch. This process took approximately 13 minutes per fold on an NVIDIA GeForce RTX 2080Ti GPU. To prevent overfitting, we implemented data augmentation on the training subset. We utilized random reflections along the horizon and vertical symmetry lines for the augmentation process: an image was reflected with a probability of 50% during each iteration.

We further augmented our data with *x* and *y* translations, where the translation distance (in pixels) is picked randomly from a uniform distribution within a chosen interval: [− 10, 10]. Lastly, we also augmented the images with random rotations of small angles between the values −5 and 5 degrees.

## 3 Results and Discussion

### 3.1 Cellular deformability analysis

To validate the connection between adhered sRBC morphology and deformability in our experimental setup, we analyzed pairs of images of the microfluidic channel first under flow (10 *µ*L/min) and then under no flow conditions. These images were examined to look for sRBCs that had not detached or moved significantly between the two image captures, to allow for legitimate comparison. The relative change in cell aspect ratio (AR) under the two flow conditions was then analyzed for each cell (Fig. 6), as a measure of cellular deformability. We have defined the cellular AR as the ratio of the estimated minor to the major axis. A set of 14 cells was identified and manually classified as seven deformable and seven non-deformable according to the morphological characteristics described in Section 1.2. After analyzing the cellular AR of the adhered RBCs under the two flow conditions, we found that the morphology of the sRBCs correlates to the deformability characteristics. The cells classified morphologically as deformable showed a mean change in AR of about 20% on average between flow and no flow. For those classified as non-deformable sRBCs, the average percent change in AR was close to zero. Results are summarized in Fig. 6. Given the heterogeneity of the cell shapes and errors introduced by the pixelation of the images, the AR changes of each subtype have a distribution, but the difference in the average AR between the two distributions is statistically significant (*p* = 0.00057). These results reproduce the link between morphology and deformability observed in Alapan *et. al*. [9] in exactly the same experimental setup we use to do the deep learning image analysis. Thus the classification into subtypes produced by the algorithm should be strongly correlated with a key biomechanical feature of the individual sRBCs.

### 3.2 Phase I network performance

#### Efficacy of class balancing using transfer learning

To test the efficacy of our approach for mitigating class imbalance in Phase I, we compared the performance of our network (pre-trained on class-balanced images) against one trained without any kind of balance control. The confusion matrix results for each network are shown in Fig. 7B, along with sample segmentation masks generated by the two networks. Since the the main utility of Phase I in our scheme is to distinguish sRBC from non-sRBC pixels, the confusion matrices are presented in terms of these two combined classes. Unsurprisingly, the imbalanced network performed poorly for the minority (sRBC) class, while our approach worked very well for both classes. This is also reflected in the trajectory of the training and validation loss for the two networks during the learning process, as shown in Fig. 4. Our network begins to outperform the imbalanced one on both the training and validation subsets early in the process, and achieves smaller losses at the end of the training.

#### Segmentation performance evaluation

To quantify overall performance of our Phase I network, we computed the performance metrics [31] defined below for a given class *i*, where *i* corresponds to either non-sRBC or sRBC pixel:

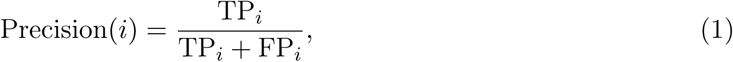

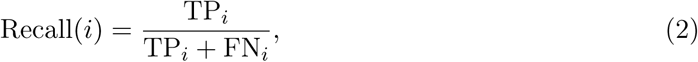

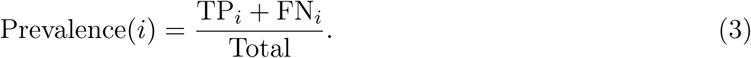

Here TP_*i*_, TN_*i*_, FP_*i*_, and FN_*i*_ denote the number of true positive, true negative, false positive and false negative outcomes in classifying a given target pixel into class *i*. “Total” represents the total number of pixels involved in the evaluation. Precision indicates the agreement between predicted and target class labels, while recall measures the the effectiveness of the neural network’s classification ability when identifying pixel-classes. Prevalence tells us how often does a specific class actually occur in the data set.

The results are summarized in Fig. 8. We see that despite the huge imbalance in non-sRBC vs. sRBC pixels (evident in the pixel count confusion matrix of Fig. 8A and the discrepancy in prevalence of Fig. 8B), our Phase I network is successful: it is able to reach state-of-the-art accuracy in segmentation of channel images from whole blood experiments compared to similar studies in literature [18, 19], reaching ≳ 97% accuracy in non-sRBC/sRBC pixel distinction in the testing set.

Care must be taken in interpreting these metrics, and they should not be naively used as an adequate standalone measure of the overall performance. As is commonly the case, we found that the bulk of our error arose from segmentation of cell boundaries rather than the cell itself. Since we are more concerned about locating centroids of the predicted segmentation masks, to crop and extract sRBC images for classification in Phase II, the cell boundary errors do not significantly affect the final results in our pipeline.

### 3.3 Phase II network performance

During learning, the network weights are optimized to make the class predictions for the training data set as accurate as possible. However, depending on the training set and the stochastic nature of the optimization process, the accuracy of the network on the testing set can vary. Attention to this issue becomes imperative when dealing with smaller data sets for classification tasks, like in our Phase II case. *k*-fold cross-validation is one approach to validate the overall performance of the network in this scenario. The general procedure starts by shuffling the total data set before splitting it into training/testing subsets, to generate an ensemble of *k* such unique subsets (or folds). We choose *k* = 5, with an 80/20% split for training/testing sets. Each fold consists of a unique combination of 20% of the images as the hold-out (test) set, and the remaining 80% as the training set. Our combined data set of 6863 total images thus generates five unique folds with training and testing sets containing 5488 and 1372 images each (accounting for rounding off). Finally, we fit the neural network parameters on the training set and evaluate the performance on the test set for five unique runs. Then for each single run, we collect the training and testing accuracy, listed in Table 2. We also show the mean and standard deviation of all the folds, with the small standard deviation being an indicator that our training did not suffer from overfitting. Performance metrics and confusion matrices for fold 1 are shown in Fig. 9, highlighting the typical ∼ 95% accuracy in object classification on the testing set.

**Table 2:** Results from the 5-fold cross-validation of the Phase II network.

| Fold No. | Training Accuracy (%) | Testing Accuracy (%) | Training Time |
| --- | --- | --- | --- |
| 1 | 97.78 | 95.63 | 13 min 25 sec |
| 2 | 98.07 | 95.39 | 13 min 47 sec |
| 3 | 98.07 | 93.59 | 14 min 36 sec |
| 4 | 97.85 | 94.75 | 15 min 18 sec |
| 5 | 97.85 | 94.97 | 15 min 10 sec |
| Mean | 97.92±0.44 | 94.77±0.74 | 14 min 45 sec |

### 3.4 Processing Pipeline: Manual vs AI Performance

After both Phase I and II are complete, we are ready for the final test of our processing pipeline, pitting the artificial intelligence (AI) deep learning approach against 3 human experts in detecting, classifying, and counting adhered sRBCs for a set of 19 whole channel images displaying a wide variety of cells. Importantly, this competition between the two sides will highlight the workflow’s true effectiveness and robustness for potentially replacing manual labor within biophysical studies of SCD. Results are illustrated in Fig. 10. Error bars along the AI axis are obtained from recall metrics of our classifier. Error bars on the manual axis are estimated from variance in repeated manual analyses on a set of whole channel images. Panel **A** shows results for total adhered sRBC cell count in each image, which can be taken as a proxy for overall object identification accuracy. We see how a very high degree of agreement is reached between our AI and human experts, with an estimated *R*^2^ statistic value of 0.96. Note how the manual error bars increase with sample size. This has serious implications for manual analyses of high cell count samples. A host of factors like longer duration of analysis time, mental fatigue of the experimentalist, etc. can affect these numbers. An AI-based automated classifier is immune to these human limitations. Panel **B** and **C** show comparison results for subcounts of deformable and non-deformable sRBCs in each sample image, indicative of classification accuracy. Excellent agreement is reached for deformable cells, with corresponding *R*^2^ ∼ 0.95. For non-deformable cells—a category significantly harder to identify because of the high degree of cross-correlating features with several objects in our “others” category—decent AI-manual agreement is still achieved, with *R*^2^ ∼ 0.77. We that note that our human experts showed significant variance among themselves in categorizing these kinds of images. The SCD pathogenetic pathway exhibits a continuum of physiological features, which we are trying to sort into discrete categories. Thus some level of disagreement is expected, particularly for images with borderline features between classes (e.g. Fig. 1B(iii) and C(i)). The role of the AI then goes beyond simply matching human performance. It takes on the role of a standardized arbiter for these borderline images, and improves overall consistency in results compared to human performance, where random, spur-of-the-moment classification decisions would play a sizeable role.

**Fig 10:**
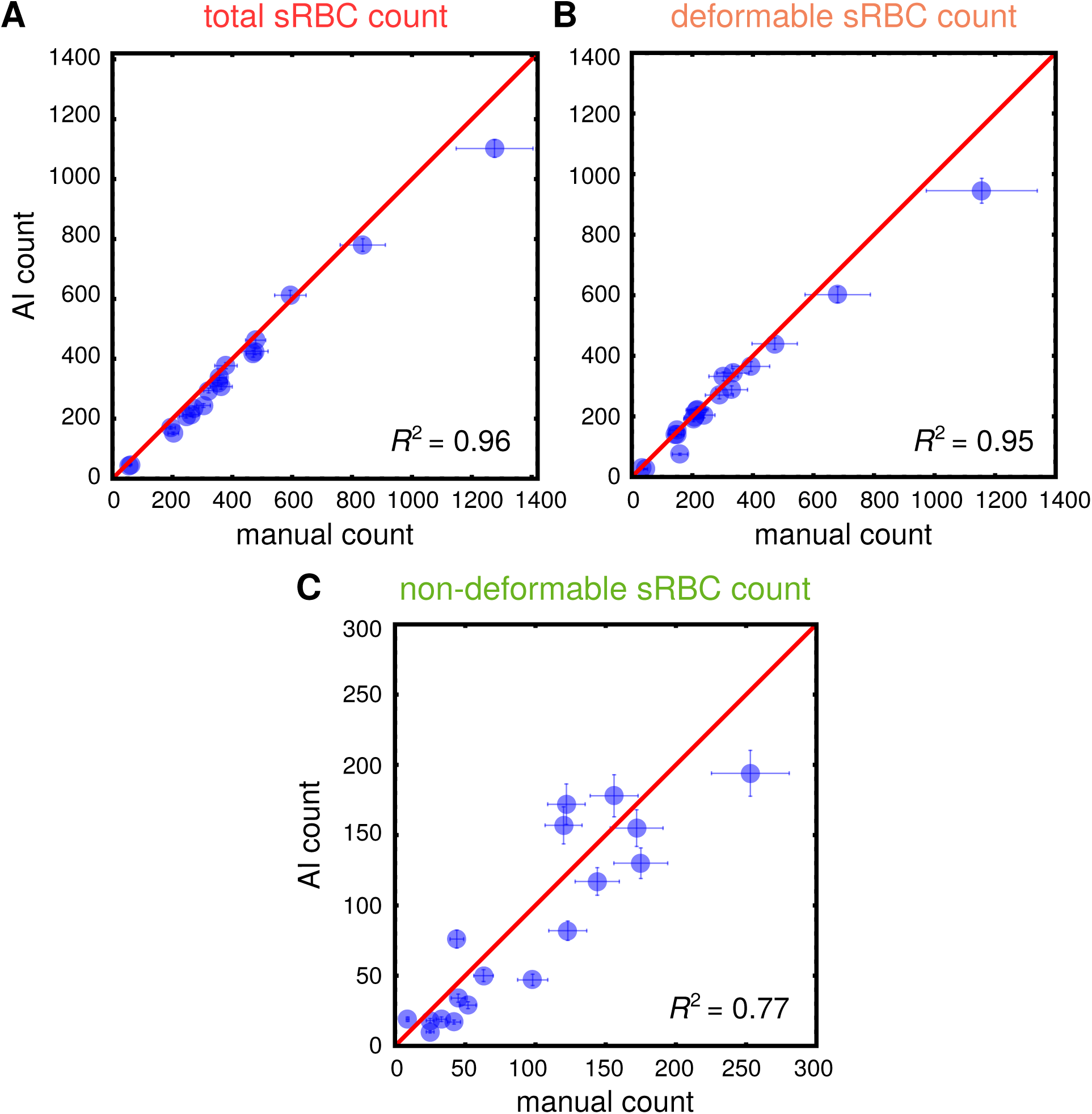
Manual vs AI performance: Results from pitting count estimates from 19 whole microchannel images processed through our automated two-part processing pipeline vs. manual characterization. Error bars along the manual axis are obtained from variance in repeated manual counts on a set of test images. The red line is the line of perfect agreement. Error bars on AI counts are estimated from the precision rates reached by our Phase II classifier network in predicting true positive outcomes in relevant categories on a test set (see Fig.9). *R*^2^ statistic values, indicating goodness of agreement between manual and AI counts, are indicated in each graph. **A:** Results for total sRBC (deformable + nondeformable) cell counts. This plot is illustrative of the high degree of accuracy achieved by our AI in identifying sRBCs. **B** and **C:** Results for number of sRBCs in each channel image classified manually and by AI as deformable or non deformable respectively. This measures the agreement reached in classification of the two morphological categories.

## 4 Conclusion

We designed and tested a deep learning image analysis workflow for a microfluidic SCD adhesion assay that completely eliminates the need for user input requiring ad hoc expertise, enabling the processing of large amounts of image data with robust and highly accurate results. For this study our target problem was identifying sRBCs in complex, whole channel bright field images using clinical whole blood samples, and distinguishing between their morphological subtypes (deformable and non-deformable). These subtypes are in turn strongly correlated with sRBC biomechanical properties, making the image analysis method a fast, high-throughput proxy for the much more laborious cell-by-cell direct measurement of membrane deformability. We demonstrated that our network performed robustly in terms of accuracy when pitted against trained personnel, while improving analysis times by two orders of magnitude.

This proof-of-concept study focuses on sRBC deformable and non-deformable classification, but this is by no means the only feature worth exploring. We are working on generalizing our workflow to examine patient heterogeneities along more expanded metrics like white blood cell (WBC) content, WBC versus sRBC content, emergent sub-types and so on. Clinical heterogeneities among SCD-affected patients constitute a critical barrier to progress in treating the disease, and understanding them will be crucial in designing targeted patient-specific curative therapies. Increasing the frequency of therapeutic interventions, novel screening technologies, and better management of both acute short term and chronic SCD complications have gone a long away in increasing patient survival. The challenge lies in achieving targeted patient-specific administration of appropriate therapies, due to the wide heterogeneity among clinical sub-phenotypes. Developing tools for consistent and comprehensive monitoring of heterogeneities among patient groups is thus paramount. Emerging potentially curative therapies like allogenic hematopoietic stem cell transplantation (HSCT) and targeted gene therapy are also promising and becoming more widely available, but mostly in developed countries. These treatments need to be streamlined and heavily standardized, requiring fast and affordable monitoring tools for assessment of efficacy checkpoints and endpoint outcomes along multiple relevant metrics. The advent of AI in high throughput, fully automated analyses of large amounts of data is able to address both these needs. The workflow presented here has been designed for integration with the SCD BioChip microfluidic assay, to ultimately realize the goal of delivering an SCD monitoring platform capable of high throughput, highly standardized and reproducible automated batch analyses enabled by AI.

## Code availability

The code associated with this manuscript is available at: https://github.com/hincz-lab/DeepLearning-SCDBiochip.

## Acknowledgments

This work was supported by the Clinical and Translational Science Collaborative of Cleveland, UL1TR002548 from the National Center for Advancing Translational Sciences component of the National Institutes of Health (NIH) and NIH Roadmap for Medical Research, Case-Coulter Translational Research Partnership Program, National Heart, Lung, and Blood Institute R01HL133574 and OT2HL152643, and National Science Foundation CAREER Awards 1552782 and 1651560. The authors acknowledge with gratitude the contributions of patients and clinicians at Seidman Cancer Center (University Hospitals, Cleveland).

